# Detecting shared independent selection

**DOI:** 10.1101/2020.04.21.053959

**Authors:** Nathan S. Harris, Alan R. Rogers

## Abstract

Signals of selection are not often shared between populations. When a mutual signal is detected, it is often not known if selection occurred before or after populations split. Here we develop a method to detect genomic regions at which selection has favored different haplotypes in two populations. This method is verified through simulations and tested on small regions of the genome. This method was then expanded to scan the phase 3 genomes of the 1000 Genomes Project populations for regions in which the evidence for independent selection is strongest. We identify several genes which likely underwent selection independently in different populations.

## 1 INTRODUCTION AND BACKGROUND

Signals of selection are sometimes shared between closely related populations (Johnson and Voight, 2018; Pickrell et al., 2009). Some of these shared signals reflect “ancestral selection,” which occurred in the population ancestral to the two populations that share the signal. Conversely, populations with similar environmental conditions may experience “independent selection,” for mutations that arose independently in the same region of the genome. Closely related populations should share more of these independent signals as well because they are more likely to live in similar environments.

However, efforts to differentiate between these two scenarios are limited. Because we cannot always identify the variant being favored, efforts to study shared signals have focused on overlapping signals (e.g. (Johnson and Voight, 2018)). More recently, Harris and DeGiorgio (2019) developed a method to distinguish between ancestral and independent selection based on measuring the difference in the frequency of sweeping haplotypes. Here we develop a method to distinguish between ancestral and independent selection without the need to determine the underlying sweeping haplotypes.

Voight et al. (2006) introduced the integrated haplotype score (iHS) to measure classic selective in genome-wide data. iHS measures the disparity in linkage disequilibrium between carriers of opposite alleles at a given site. Large negative iHS values indicate a disproportionate amount of LD around the derived allele, implying that it is has increased in frequency relatively rapidly. Large positive values of iHS indicate a similar scenario for the ancestral allele. Usually, only the magnitude of iHS is considered, as new beneficial alleles, the target of selection, occur on haplotypes with an essentially random arrangement of ancestral and derived alleles.

Retaining the sign of iHS provides information about variation on the favored haplotype. While two populations may have similar |iHS| magnitudes, scores at individual sites may have opposite signs. This situation indicates selection at a site in both populations, but for different alleles. This will likely happen when two populations split before selection occurred and the background variation around independently selected loci reflects independently accumulated variation in each lineage. If two populations have split recently, a beneficial mutation sweeping in their common ancestor may end up in both daughter populations, along with the haplotype on which it occurred. In this scenario, two populations would likely have similar iHS magnitudes *and* signs.

## 2 RESULTS

### The independent selection index

Comparing iHS values while retaining the sign allows indirect comparison of sweeping haplotypes. We use this principal to develop a method for identifying genomic regions in which positive selection has occurred independently in two populations. Within 100-kb windows, we calculated the Independent Selection Index (ISI):

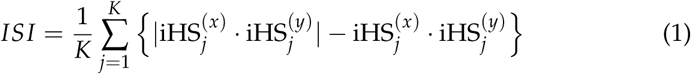

where *j* indexes the *K* sites within the window, and iHS_*j*_^(*z*)^ is the signed iHS value at site *j* in population *z*. The *j*th term in this sum equals zero when iHS has the same sign in both populations but is positive if the signs differ. *ISI* will be near zero when the same haplotype has been favored in both populations, because in that case, the signs will be the same. *ISI* becomes increasingly positive when different haplotypes are favored, because then the signs will tend to differ.

Simulations were performed using Selection on Linked Mutations (SLiM) package (Messer, 2013; Haller and Messer, 2018). Simulations were run using three scenarios: neutral, positive selection before population splits, and positive selection following population splits. In each case, simulations modelled a single population that splits in two at a range of pre-specified times. Neutral scenarios contain only mutations with no effect. In both cases of positive selection, a single beneficial mutation is introduced into the population. In half of the simulations a functionally equivalent mutation occurs in both populations at the same site following the population split. This represents the scenario in which two populations experience independent selection at the same locus following the split from a common ancestor. The placement of these mutations ensures that signals of selection will be in the same location in the simulated data, and the similarity of iHS values of sites around the introduced mutations will affect ISI. In the other half of the positive simulations, the beneficial allele is introduced in the common ancestor of the two populations. Beneficial mutations arise in the ancestral population during an interval (*t*, 2*t*), measured backwards from the present. Here, *t* is the time when the ancestral population splits. The mutations remain advantageous, even after the split. On average, these mutations are under positive selection twice as long as in the previous model. Because of this, the selective advantage in these scenarios is halved.

In all simulations a single beneficial mutation occurs in the middle of the chromosome. Large ISI values should occur where both populations have large iHS scores with opposite signs at the same loci. Figure 1 shows the simulation with the largest value of ISI. Figures 2 and 3 show Manhatten plots for ISI for the different divergence times. We find that the simulations in which the beneficial mutation is under selection in the common ancestor of two populations do not produce extreme values of this statistic, and the largest scores are randomly distributed across the simulated chromosome (Figure 4). Large values of ISI occur when selection has occurred independently following the split from the common ancestor. The regions with the largest ISI values in these cases either contain the causative variant, or are adjacent to it.

**Figure 1:**
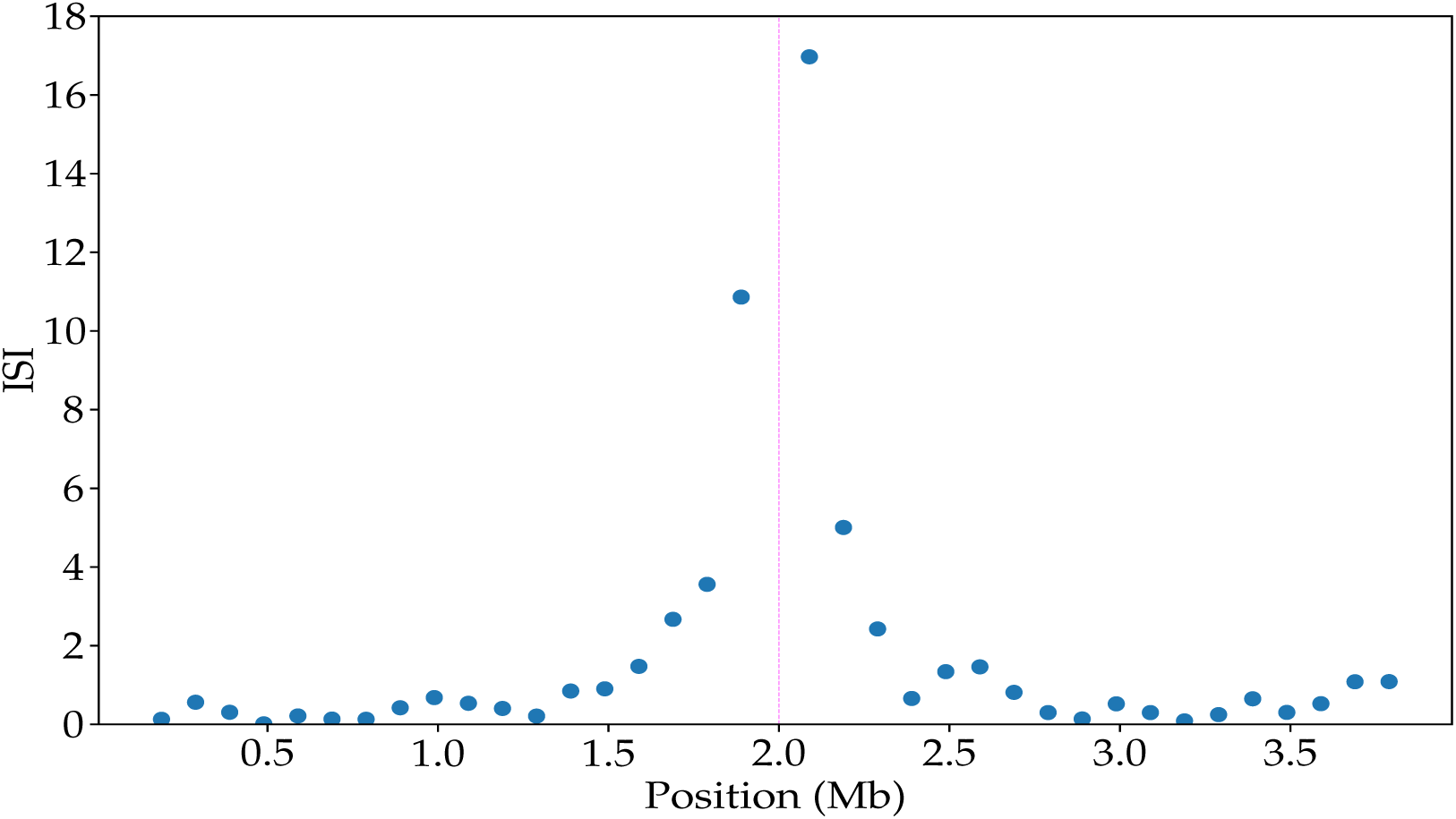
For illustration, the results of the simulation with the largest value of ISI. ISI between simulated populations for simulations in which a single beneficial mutation occurs *after* a population split.

**Figure 2:**
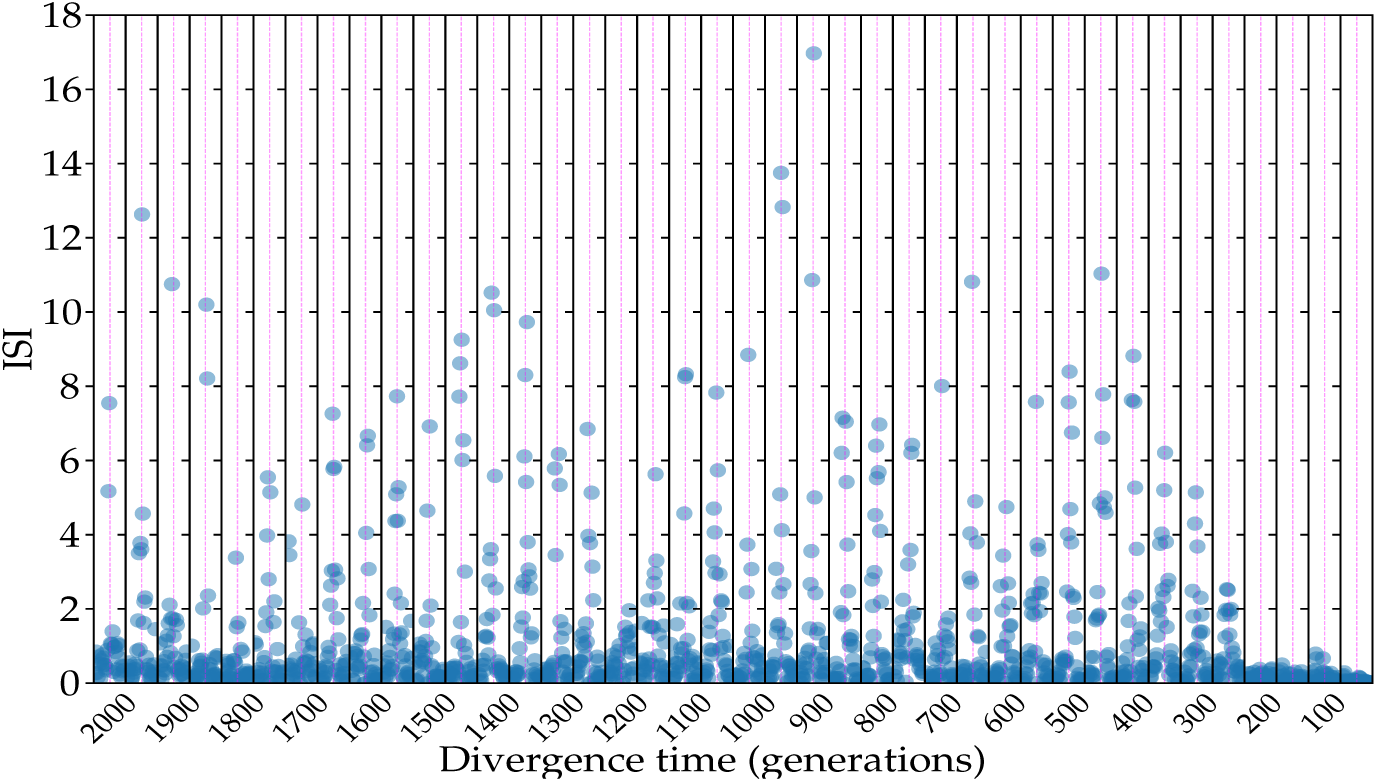
ISI between simulated populations for simulations in which a single beneficial mutation occurs *after* a population split. With the exception of very recent divergence times, the signal of independent selection is identified. The beneficial mutation is placed in the middle of the chromosome, indicated by the dashed line.

**Figure 3:**
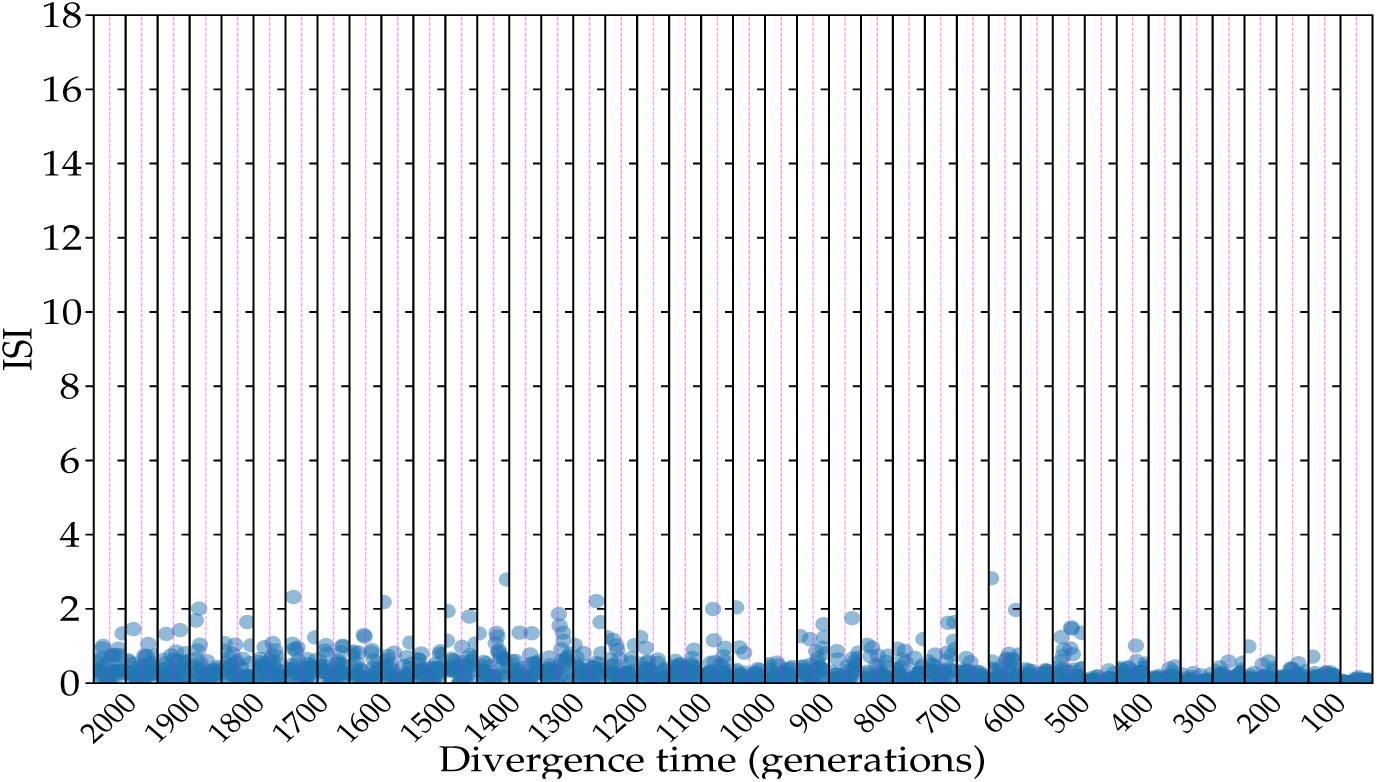
ISI between simulated populations for simulations in which a single beneficial mutation occurs *before* a population split. Both populations experience a signal of selection in the same region, but it is not detected at any divergence time. The beneficial mutation is placed in the middle of the chromosome, indicated by the dashed line.

**Figure 4:**
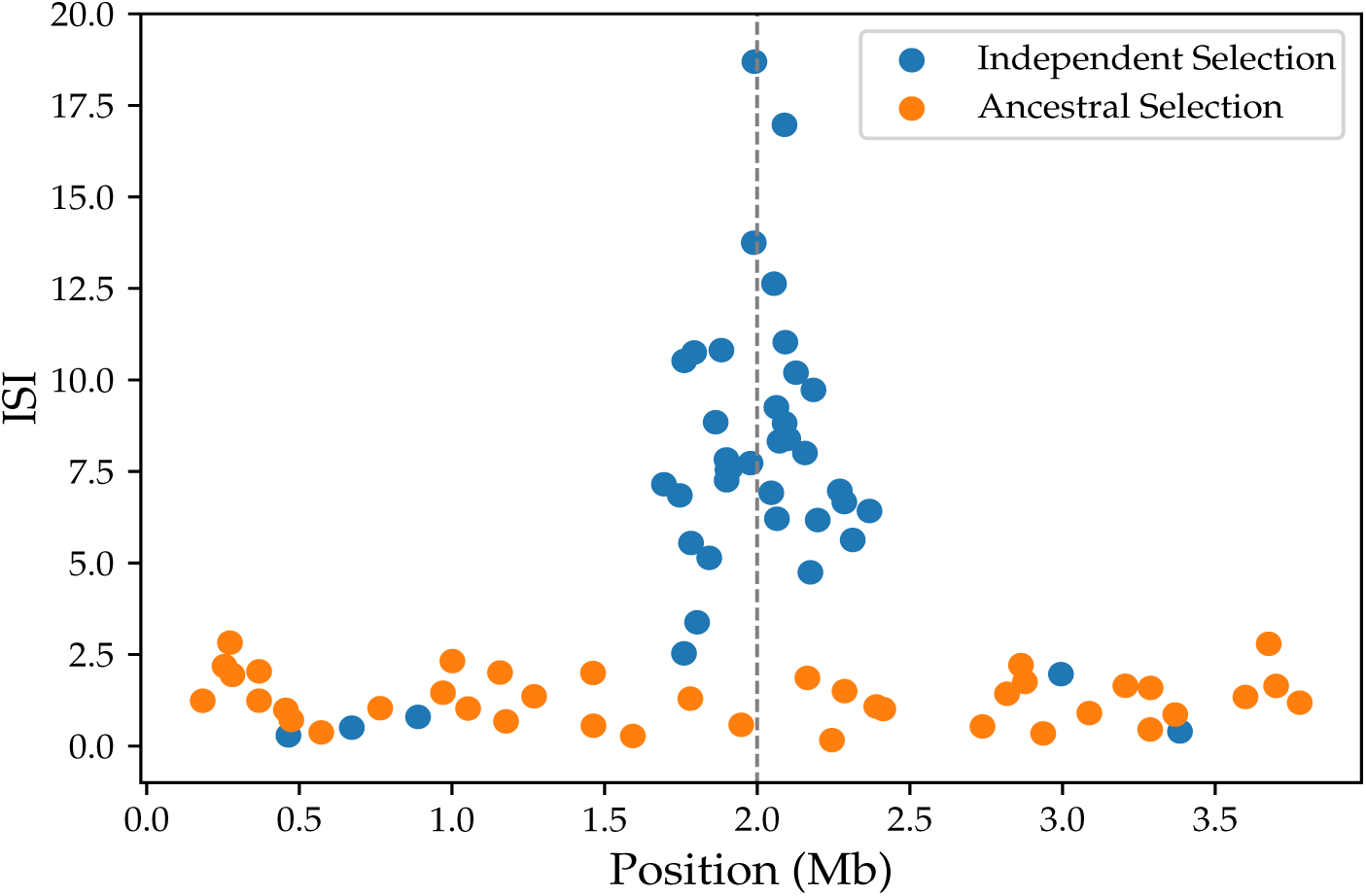
The top ISI scores from each simulation are plotted together. In general, we see that ISI successfully identifies regions near the introduced beneficial mutation (vertical dashed line) when selection occurs after a population split bot not when it occurs before. There are five divergence times at which ISI produces a false negatives. Four of these are the most recent divergence times, suggesting ISI is not sensitive to independent selection in the very recent past.

### Cases of independent Selection

Next, this method was applied to LCT and the glycophorin cluster. Figure 5 shows ISI values around the LCT gene in European populations. Not surprisingly, most population pairs have very low values of ISI. However, comparisons with TSI have relatively elevated ISI values, implying that selection at LCT may be occurring on different haplotypes than those sweeping in the rest of Europe. The signal around the glycophorin cluster is shared across a wider range of populations, with large iHS signals present in all 1000 genomes populations measured (see Appendix C.). Variation in beneficial haplotypes occurs both within continental regions (Figure 6), and between continental regions (Figure 7) as predicted above.

**Figure 5:**
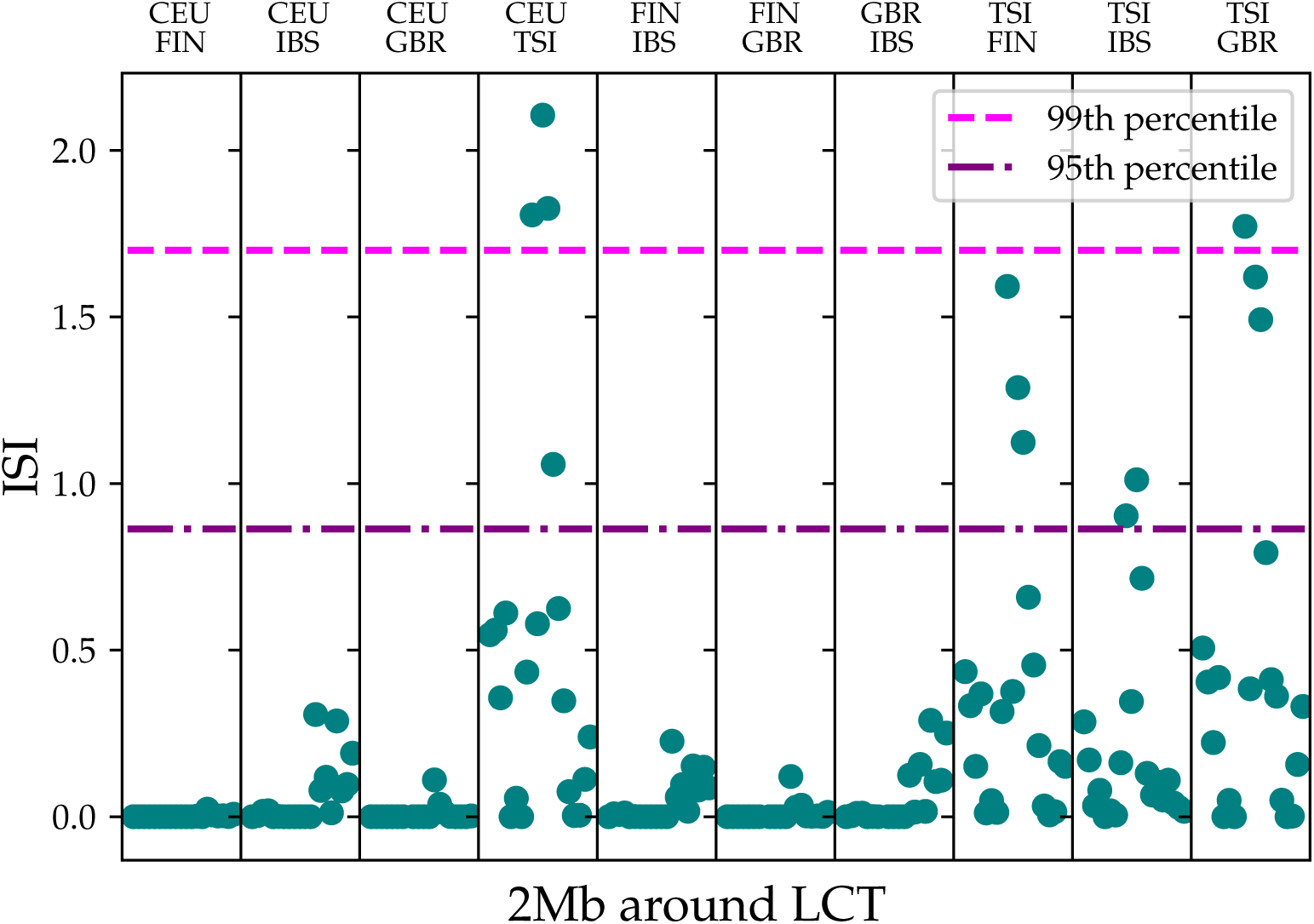
ISI scores for the LCT region in European populations. TSI has the weakest signal of selection in individual analysis (see Appendix C), but the largest values of ISI in the lactase region occur when TSI is compared to other European populations. This suggests that the haplotype under selection in TSI is distinct from that in other European populations, and selection on lactase in Europe is probably occurring on multiple haplotypes.

**Figure 6:**
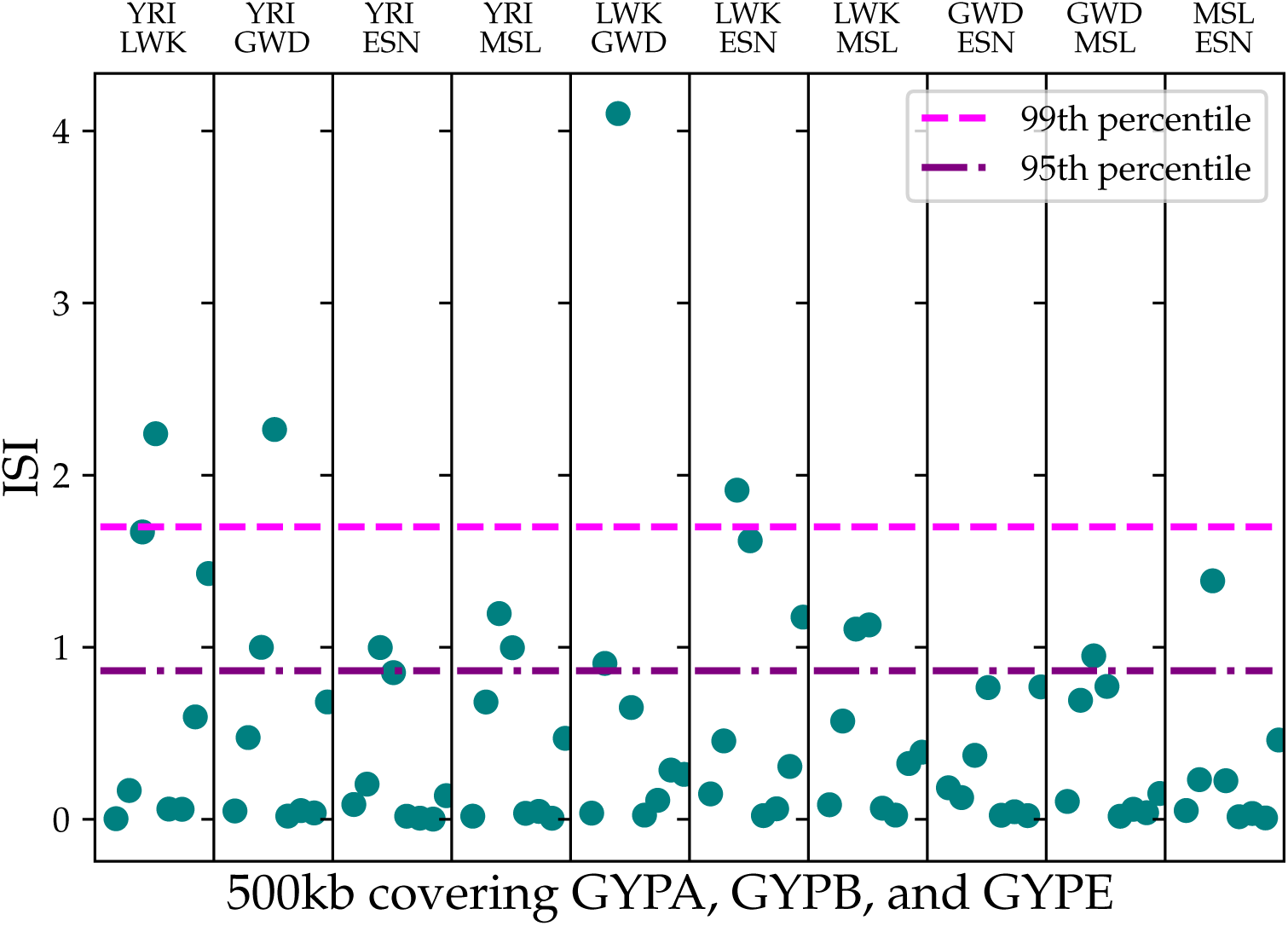
ISI between African populations around *GYPA*, *GYPB*, and *GYPE*. Both ancestral and independent selection is present within the African samples.

**Figure 7:**
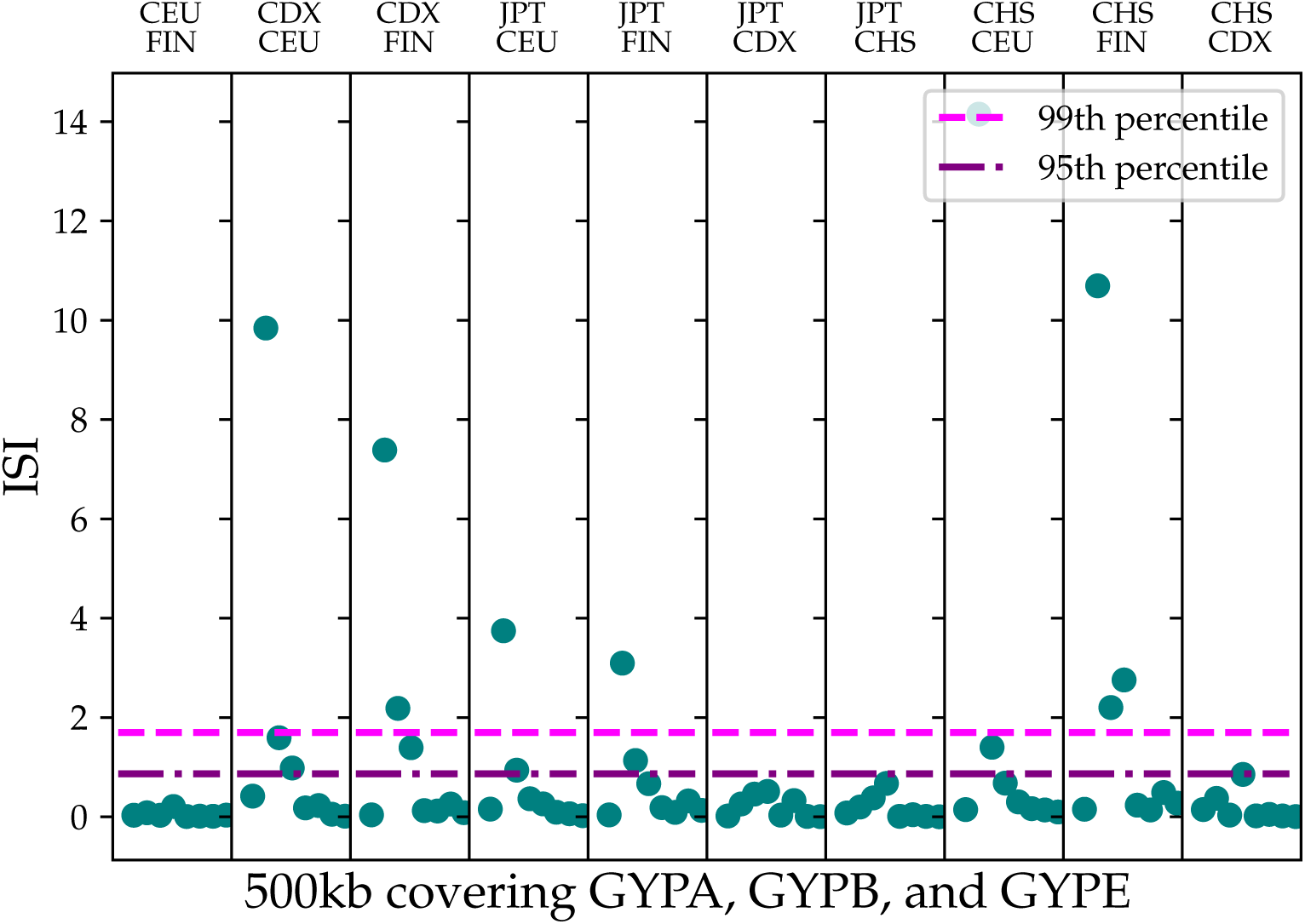
ISI between European and East Asian populations around *GYPA*, *GYPB*, and *GYPE*. Selection within continental regions appears to occur on a single haplotype, but occurring on independent haplotypes in the two continents.

### ISI across the genome

This method was next applied to every pair of 1000 Genomes Phase 3 populations across the entire genome. Figure 8 shows an example of ISI across the genome for the Vietnamese (KHV) and English (GBR) samples. Because some populations may share more independent selection than others, all potential population pairs are considered together when looking for the largest values of ISI. Table 3.1 shows the top regions returned from this analysis. These results were picked from regions with at least 50 shared SNPs between populations and ISI greater than five.

**Figure 8:**
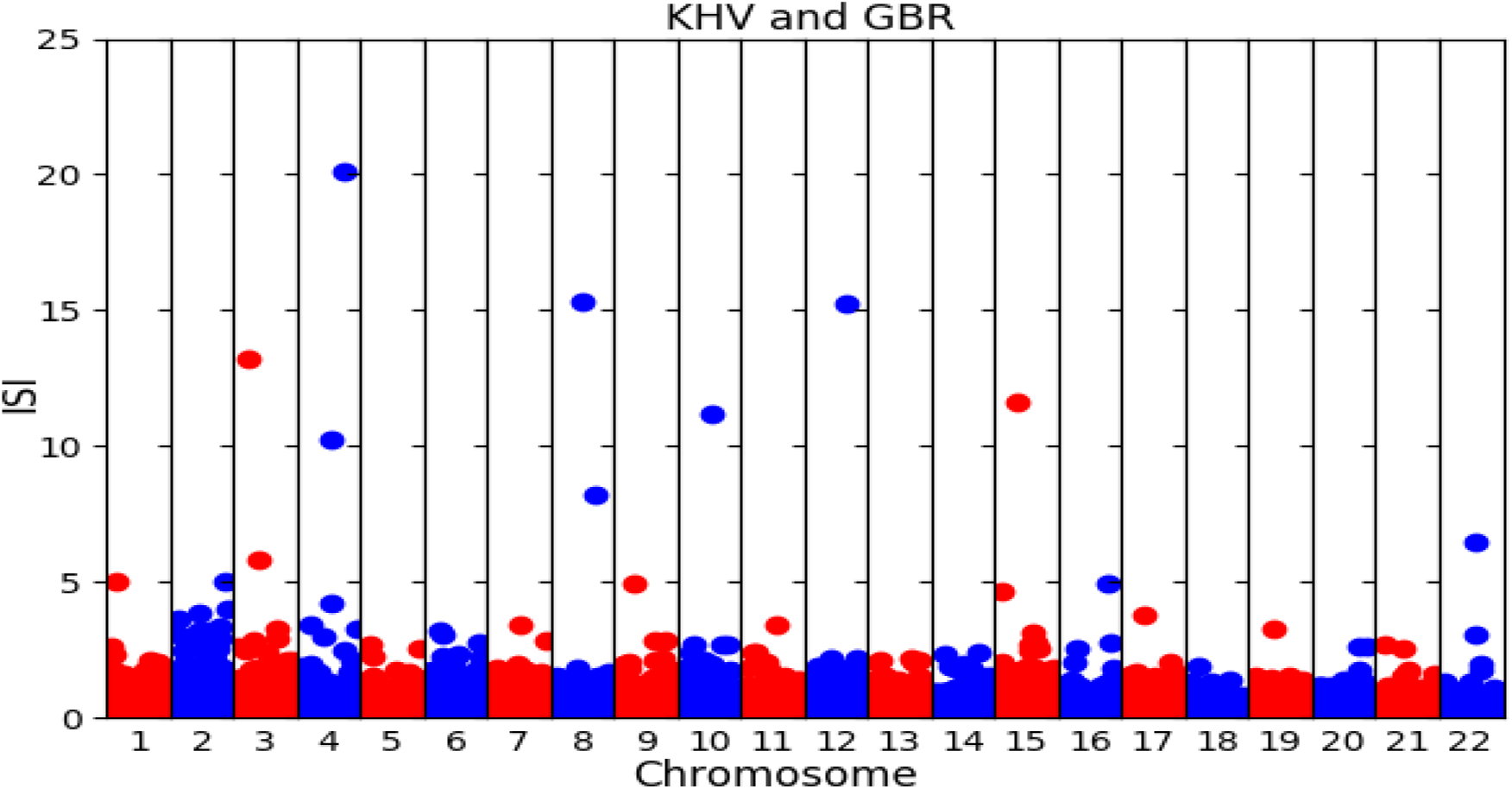
ISI plotted for each chromosome in the vietnamese(KHV) and English (GBR) pair. We find that some outliers are specific to population pairs, such as he outlier on chromosome three, while other outliers are found in many population pairs, such as the outliers on chromosome ten.

These regions were scanned for overlap with known coding regions (Table 2). Not all of the regions in Table 1 are present, as some of the results are found entirely in non-coding regions. The genes listed here are good candidates for independent selection. The function and associations of these candidates varies considerably. Examples include: Mitochondrial transporters (*SLC25A32*) (Spaan et al., 2005), issues with mitochondrial translation (*MRPS16*) (Miller et al., 2004), issues with melanoblast migration and cancer (*P-REX1*) (Lindsay et al., 2011), MNS blood group expression (*GYPE*) (Willemetz et al., 2015), vulvar cancer cell proliferation (*MIR3147*) (Yang and Guo, 2018), and neural tube defects (*FZD6*) (De Marco et al., 2012).

**Table 1:**
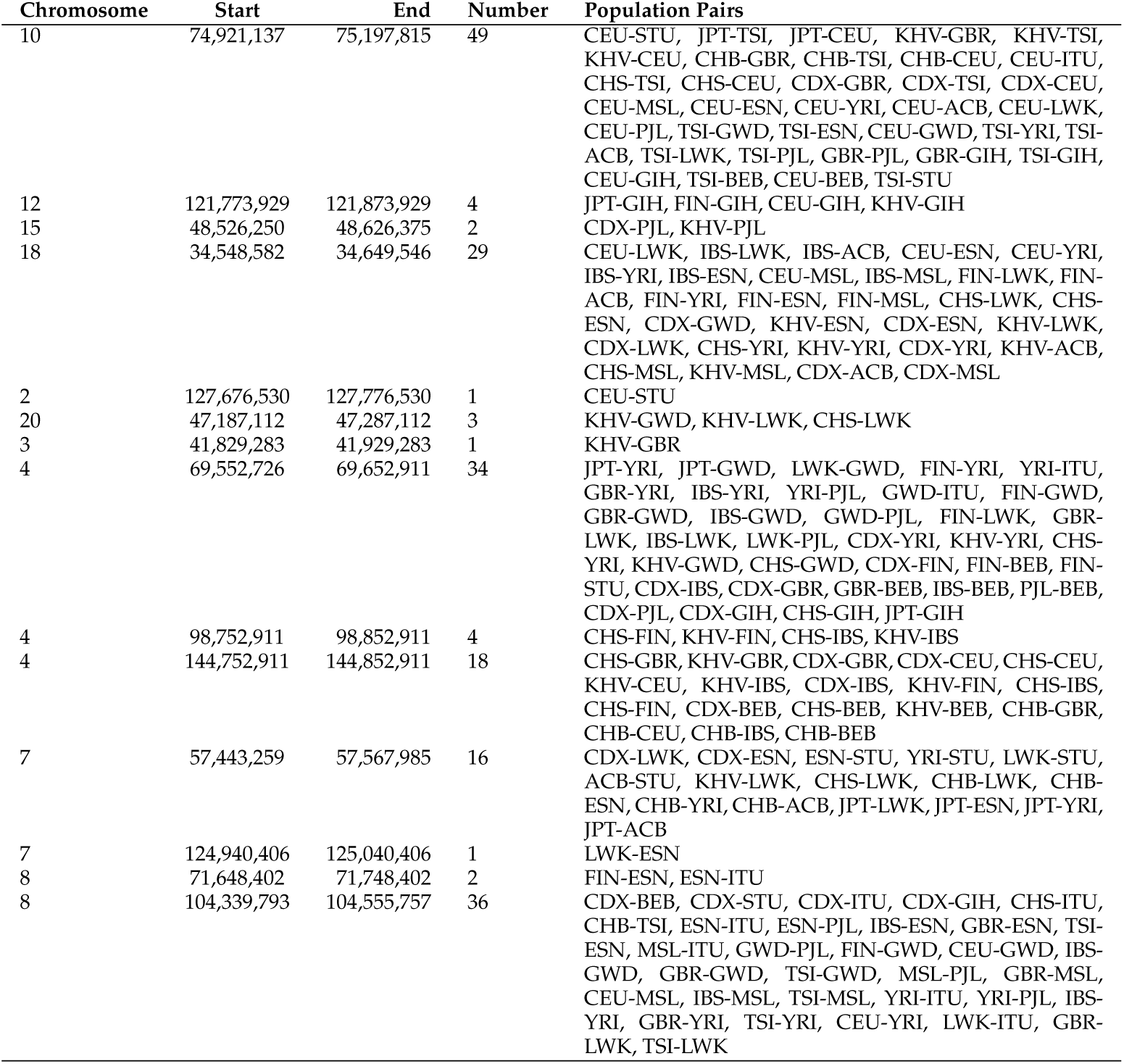
Candidate regions of the genome that show the most extreme measures of independent selection in two populations.

| Chromosome | Start | End | Number | Population Pairs |
| --- | --- | --- | --- | --- |
| 10 | 74,921,137 | 75,197,815 | 49 | CEU-STU, JPT-TSI, JPT-CEU, KHV-GBR, KHV-TSI, KHV-CEU, CHB-GBR, CHB-TSI, CHB-CEU, CEU-ITU, CHS-TSI, CHS-CEU, CDX-GBR, CDX-TSI, CDX-CEU, CEU-MSL, CEU-ESN, CEU-YRI, CEU-ACB, CEU-LWK, CEU-PJL, TSI-GWD, TSI-ESN, CEU-GWD, TSI-YRI, TSI-ACB, TSI-LWK, TSI-PJL, GBR-PJL, GBR-GIH, TSI-GIH, CEU-GIH, TSI-BEB, CEU-BEB, TSI-STU |
| 12 | 121,773,929 | 121,873,929 | 4 | JPT-GIH, FIN-GIH, CEU-GIH, KHV-GIH |
| 15 | 48,526,250 | 48,626,375 | 2 | CDX-PJL, KHV-PJL |
| 18 | 34,548,582 | 34,649,546 | 29 | CEU-LWK, IBS-LWK, IBS-ACB, CEU-ESN, CEU-YRI, IBS-YRI, IBS-ESN, CEU-MSL, IBS-MSL, FIN-LWK, FIN-ACB, FIN-YRI, FIN-ESN, FIN-MSL, CHS-LWK, CHS-ESN, CDX-GWD, KHV-ESN, CDX-ESN, KHV-LWK, CDX-LWK, CHS-YRI, KHV-YRI, CDX-YRI, KHV-ACB, CHS-MSL, KHV-MSL, CDX-ACB, CDX-MSL |
| 2 | 127,676,530 | 127,776,530 | 1 | CEU-STU |
| 20 | 47,187,112 | 47,287,112 | 3 | KHV-GWD, KHV-LWK, CHS-LWK |
| 3 | 41,829,283 | 41,929,283 | 1 | KHV-GBR |
| 4 | 69,552,726 | 69,652,911 | 34 | JPT-YRI, JPT-GWD, LWK-GWD, FIN-YRI, YRI-ITU, GBR-YRI, IBS-YRI, YRI-PJL, GWD-ITU, FIN-GWD, GBR-GWD, IBS-GWD, GWD-PJL, FIN-LWK, GBR-LWK, IBS-LWK, LWK-PJL, CDX-YRI, KHV-YRI, CHS-YRI, KHV-GWD, CHS-GWD, CDX-FIN, FIN-BEB, FIN-STU, CDX-IBS, CDX-GBR, GBR-BEB, IBS-BEB, PJL-BEB, CDX-PJL, CDX-GIH, CHS-GIH, JPT-GIH |
| 4 | 98,752,911 | 98,852,911 | 4 | CHS-FIN, KHV-FIN, CHS-IBS, KHV-IBS |
| 4 | 144,752,911 | 144,852,911 | 18 | CHS-GBR, KHV-GBR, CDX-GBR, CDX-CEU, CHS-CEU, KHV-CEU, KHV-IBS, CDX-IBS, KHV-FIN, CHS-IBS, CHS-FIN, CDX-BEB, CHS-BEB, KHV-BEB, CHB-GBR, CHB-CEU, CHB-IBS, CHB-BEB |
| 7 | 57,443,259 | 57,567,985 | 16 | CDX-LWK, CDX-ESN, ESN-STU, YRI-STU, LWK-STU, ACB-STU, KHV-LWK, CHS-LWK, CHB-LWK, CHB-ESN, CHB-YRI, CHB-ACB, JPT-LWK, JPT-ESN, JPT-YRI, JPT-ACB |
| 7 | 124,940,406 | 125,040,406 | 1 | LWK-ESN |
| 8 | 71,648,402 | 71,748,402 | 2 | FIN-ESN, ESN-ITU |
| 8 | 104,339,793 | 104,555,757 | 36 | CDX-BEB, CDX-STU, CDX-ITU, CDX-GIH, CHS-ITU, CHB-TSI, ESN-ITU, ESN-PJL, IBS-ESN, GBR-ESN, TSI-ESN, MSL-ITU, GWD-PJL, FIN-GWD, CEU-GWD, IBS-GWD, GBR-GWD, TSI-GWD, MSL-PJL, GBR-MSL, CEU-MSL, IBS-MSL, TSI-MSL, YRI-ITU, YRI-PJL, IBS-YRI, GBR-YRI, TSI-YRI, CEU-YRI, LWK-ITU, GBR-LWK, TSI-LWK |

**Table 2:** Genes that intersect with the regions of the genome that show the most extreme measures of independent selection in two populations. Some regions from Table 1 are absent here because they occur in non-coding regions.

| Chromosome | Start | End | Gene Symbol |
| --- | --- | --- | --- |
| chr10 | 74,921,137 | 75,189,422 | ECD, FAM149B1, DNAJC9, MRPS16, DNAJC9-AS1, ANXA7, MSS51, CFAP70 |
| chr12 | 121,773,929 | 121,873,929 | ANAPC5, BC029038, RNF34, KDM2B |
| chr15 | 48,526,250 | 48,626,375 | SLC12A1, DUT |
| chr18 | 34,548,582 | 34,649,546 | KIAA1328 |
| chr20 | 47,240,792 | 47,287,112 | AX746653, PREX1 |
| chr3 | 41,829,283 | 41,929,283 | ULK4 |
| chr4 | 98,752,911 | 98,852,911 | STPG2 |
| chr4 | 144,797,907 | 144,826,660 | GYPE |
| chr4 | 144,833,483 | 144,833,483 | BC029578 |
| chr7 | 57,472,730 | 57,472,730 | MIR3147 |
| chr7 | 57,476,012 | 57,476,012 | DQ578920 |
| chr7 | 57,509,994 | 57,529,655 | ZNF716 |
| chr8 | 104,339,793 | 104,343,737 | FZD6 |
| chr8 | 104,383,884 | 104,394,828 | CTHRC1 |
| chr8 | 104,412,638 | 104,427,165 | SLC25A32 |
| chr8 | 104,427,218 | 104,455,110 | DCAF13 |
| chr8 | 104,513,114 | 104,555,757 | RIMS2 |

## 3 DISCUSSION

### Using the sign of iHS

Results from the simulations show that whether selection occurs before or after a population split affects the correlation of signed iHS, but not unsigned iHS. This result is useful because it allows discrimination between independent selection and selection in a common ancestor for a particular region of interest.

Lactase is known to have been under recent selection in several geographic regions (Ségurel and Bon, 2017), but the genetic cause is thought to have a single origin in Europe. The results from iHS support this previous finding. Not only do four out of the five populations in Europe show evidence for shared selection in the LCT region, the correlation of signed iHS implies the variation around the selected variant is consistent with a single beneficial haplotype. However, the fifth population, the Toscani, shows significant values of ISI around LCT. This suggests that selection around LCT in Europe was more diverse than previously believed. There is some evidence in the literature that the Eurasian genotype has arisen on more than one haplotype independently (Enattah et al., 2007). For TSI specifically, Schlebusch et al. (2013) found that a sweeping lactase persistence haplotype in the Maasai from Kenya was three times as common in Tuscans as the European haplotype. However, the variant associated with lactase persistence in the Maasai, while present in the 1000 Genomes population from Kenya, is missing from the TSI sample. So while we can confirm the presence of selection around LCT in the Toscani, we cannot conclude that the same variant is sweeping.

The glycophorin cluster has primarily been associated with malarial resistance in African populations (Wang et al., 2003). The presence of signals around these genes in populations outside the malarial zone suggests independent selective pressure on immune characteristics at these loci, supporting previous work (Bigham et al., 2018). The most striking result is the presence of shared signals in both Asian and European populations, in which the beneficial haplotype is distinct not only from one another, but from the African haplotype as well. These results therefore support at least three independent origins of selection around the glycophorin cluster, two of which are unlikely to be driven by malaria. A table containing ISI values for all population comparisons can be found in Appendix C.

Selected haplotypes also vary within geographic regions. ISI scores within Africa vary from small values, like between ESN and GWD (0.525) to large ISI values between LWK and GWD (5.65), the former implying selection on a shared haplotype and the latter implying selection on independent haplotypes. However, most values within Africa are intermediate in size, falling under, but close to, the cutoff of 1.459 for the top one percent of ISI scores. Considering that large iHS scores are present in each population, the intermediate values imply that some selection in this region is shared among African populations, while some is not. This is consistent with the observation that Africans have a larger number of variants in the glycophorin cluster known to be associated with disease (Leffler et al., 2017), allowing simultaneous selection of variants in multiple exons rather than a single beneficial haplotype. In contrast, selection in this region in European populations can largely be traced to a single beneficial haplotype. ISI values are small for pairwise comparisons in Europe with the exception of the Toscani, whose smallest ISI score occurs when compared with the South Asian population GIH, or Gujarati Indians from Houston, Texas. This exception to the pattern may be caused by the introduction of a beneficial haplotype from one population to another, or into each from a third population, but further work will need to be done to investigate such population specific examples.

While there are some population pairs with significant ISI values around well documented genes like LCT, the most extreme values of ISI are found elsewhere. The results of the genomic ISI selection scan provide a finer view of shared selection. In some cases, relatively few population pairs display evidence for independent selection at a locus. For example, chromosome three contains a candidate region that is shared only by the Vietnamese (KHV) and the English (GBR) (Table 2). However, there also seems to be independent selection occurring at the level of continental groups rather than individual populations. For example, the candidate region on chromosome 10 had the largest value of ISI, but is also found in many population pairs. Furthermore, the population pairs have a seemingly non-random pattern. This candidate region shows up in comparisons between Europeans and Africans, Asians and Africans, and Europeans and Asians. However, it does not appear when any of the populations within each geographic region are compared to one another. These patterns can be seen visually by comparing the direction of the iHS scores in Figure 9. A possible explanation for this pattern is independent instances of selection in the same region of the genome, and a small sample size in the African and South Asian comparison meant it was filtered out. Another possibility is that South Asians and Africans actually share more of their haplotype.

**Figure 9:**
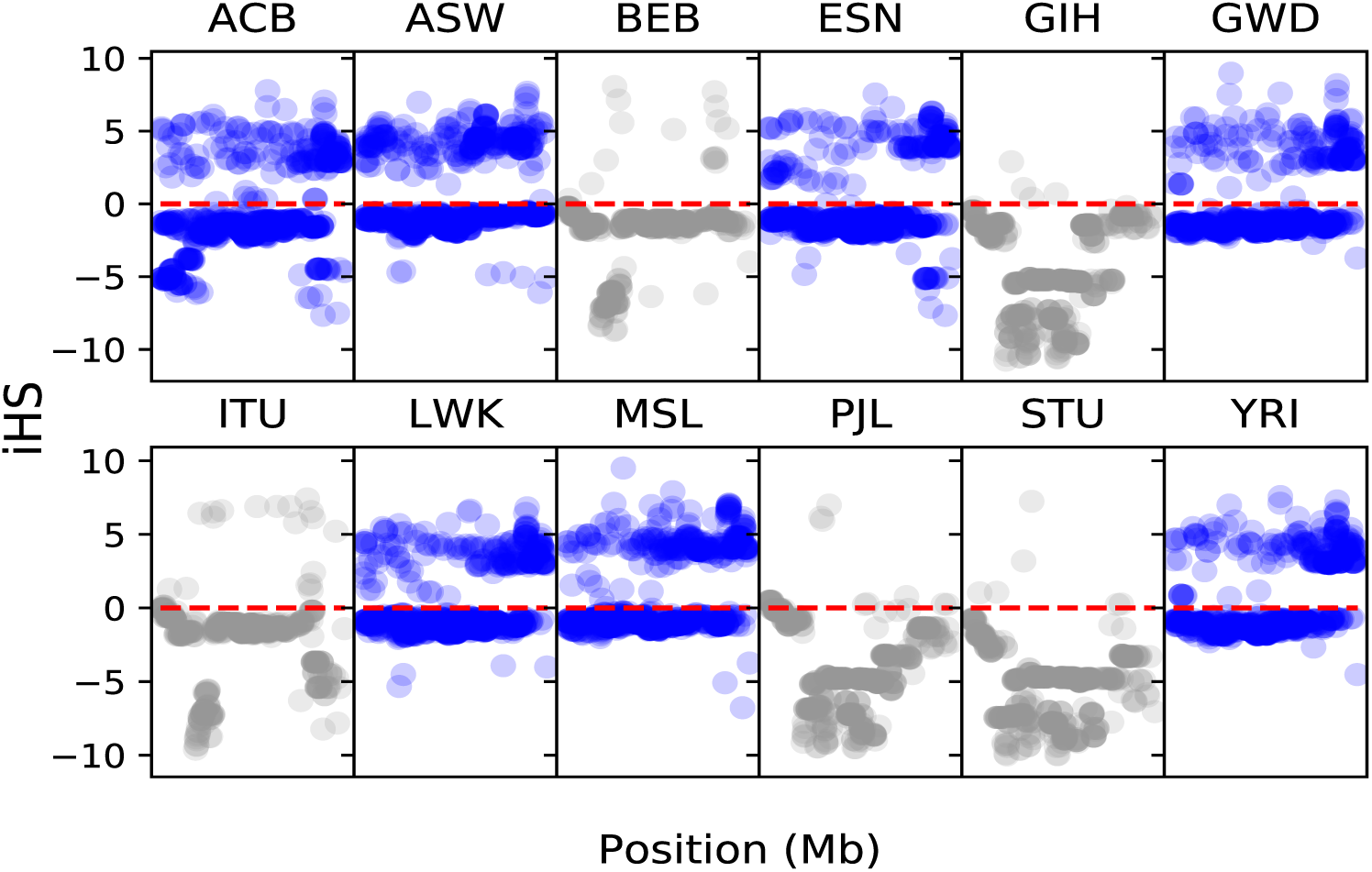
iHS results for candidate region located on chromosome 10 in African (blue) and South Asian (gray) populations.

The methods developed here allow new insights into well studied loci such as LCT, and have the potential to scan for new signals of loci under selection by considering the sign of iHS. Specifically, ISI shows that patterns of shared independent selection may occur for specific population pairs or between geographic regions. Instances of shared selection will need to be investigated individually and we are optimistic that the work presented here will open new avenues of research.

## 4 METHODS

### Data

The 1000 Genomes Phase 3 variant data were obtained from ftp://ftp.1000genomes.ebi.ac.uk/vol1/ftp/ for populations with ancestry from South Asia, East Asia, Europe, and Africa. American populations and the African American sample were removed due to recent extensive admixture. Multiallelic sites were removed.

### Correlation of iHS and correlation of |iHS|

iHS and |iHS| were calculated using the *Selscan* package (Szpiech and Hernandez, 2014) for each population in the 1000 Genomes Phase 3 data. Correlation of iHS (signed) scores from one population with scores from another was calculated for each possible pair of populations. This process was repeated for |iHS|. Confidence intervals were generated using a moving blocks bootstrap (Liu and Singh, 1992) with a block size of 500 kb.

The LCT and glycophorin cluster subdivisions contained a megabase of flanking region and 125kb flanking regions respectively. The difference in the choice of flank size reflects the size of the regions. A flanking region around LCT was used to increase this sample size and allow bootstrap replicates to be used to generate confidence intervals. This makes sense in the case of LCT because the block of LD surrounding the locus is exceptionally large because selection at LCT was especially strong and relatively recent. The small flanking region for the glycophorin cluster was used to increase the size of the region to make the bootstrap replicate size used for the genome as a whole. While this region is still too small for bootstrap replicates, this approach allows us to directly compare the resulting 500kb region to genic or nongenic regions in the genome.

### Simulations

Simulated data were generated using the Selection on Linked Mutations (SLiM) package (Messer, 2013; Haller and Messer, 2018). In each simulation a single population splits into two at a pre-specified time. A set of neutral simulations were run first to generate a set of neutral data to compare to models that include selection.

Two types of simulations with selection were conducted. First, a single beneficial mutation (*N* = 10, 000 *s* = 0.025) is introduced into the population before it splits. If this mutation is not lost to drift, the simulation continues with selection until the present. In the second model, a beneficial mutation (*N* = 10, 000, *s* = 0.05) is inserted into both populations following the split. These mutations are inserted at the same location in the middle of the chromosome, and the simulation is restarted if either is lost to drift. The difference in selection coefficient between the two models exists because we wanted to test the effect of a beneficial allele that started in an ancestral population but can continue to sweep in daughter populations. This being the case, selection will be occur for twice as long in the models in which the beneficial allele is introduced before the population split. However, we did not want the results of these two models to differ due to length of selection. We therefore halved the selection coefficient in the ancestral-selection model, because the time required for any given change is proportional to 1/*s* (Crow et al., 1970).

iHS was calculated for each simulation using *selscan* (Szpiech and Hernandez, 2014). Due to the sensitivity of iHS to small allele frequencies, sites with a minor allele frequency less than 0.05 were removed. Simulations with selection were standardized for allele frequency jointly with neutral simulations with the same divergence time. We standardized each iHS value by subtracting off the mean iHS across the genome for sites in the same allele-frequency bin. These means were calculated within 10 bins of allele frequency, spanning the range from 0 to 1.

### Candidate regions for independent selection

To find candidate regions for independent selection, each population pair was divided into 100 kb regions. ISI was calculated for each region. ISI was favored over covariance and correlation because of differences in variance between the populations. In addition, the statistic is intuitive, as the two terms it contains should approach the same value when iHS scores have the same sign.

The results from all population pairs were concatenated and the results with the top one percent of ISI were observed. The regions with the most extreme scores commonly had very small sample sizes. As a result, we trimmed the results to only include regions with at least 50 SNPs shared between the population pair. All population pairs were considered together because there is no reason to suspect that the number of shared signals due to independent selection is similar in each pair. For example, populations that split very recently are less likely to show evidence of independent selection because little time has elapsed and they more likely to share ancestral haplotypes.

