## Supplemental Tables and Figures for "Detecting shared independent selection"

SUPPORTING INFORMATION FOR: DETECTING SHARED INDEPENDENT SELECTION

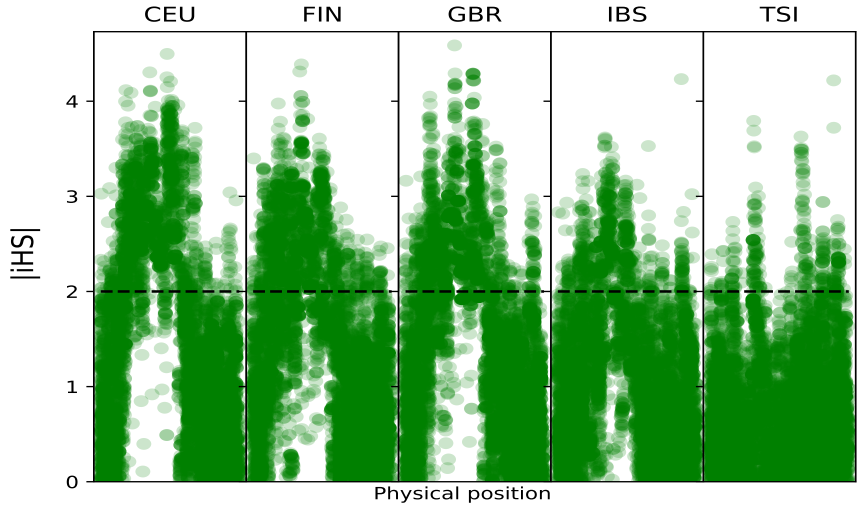


Figure 1: Mutual evidence of selection at LCT in European populations.

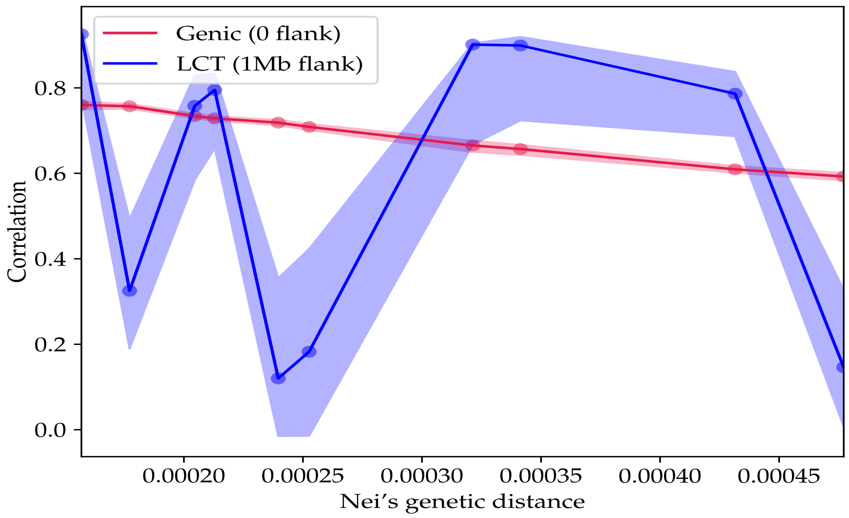


Figure 2: Correlation of |iHS| in LCT. The trend is in LCT is unclear because the Toscani (TSI) are included but do not show evidence of strong selection at LCT.

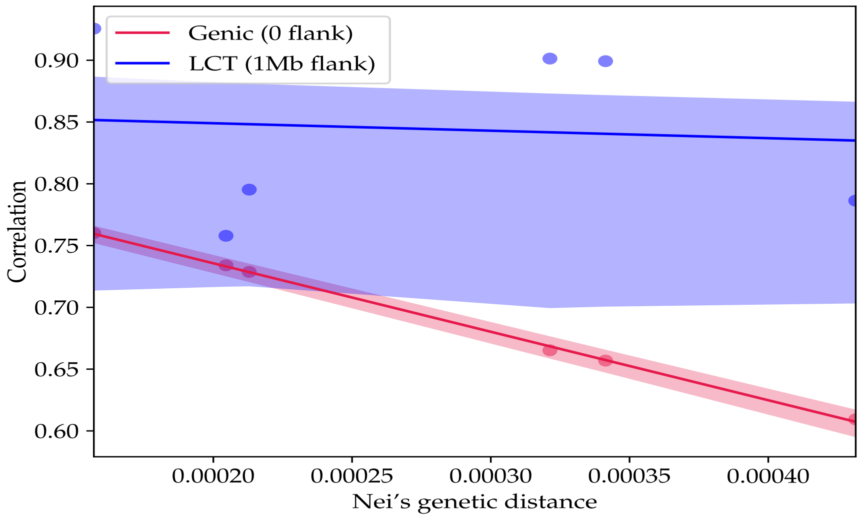


Figure 3: Correlation of |iHS| in LCT in Europe excluding the Toscani (TSI). The trend in correlation implies a strong correlation at the same locus in the remaining European populations.

Table 1: 1000 Genomes Phase 3 population codes.

| Population Code | Population Details | Super Population Code |
|---|---|---|
| CHB | Han Chinese in Beijing, China | EAS |
| JPT | Japanese in Tokyo, Japan | EAS |
| CHS | Southern Han Chinese | EAS |
| CDX | Chinese Dai in Xishuangbanna, China | EAS |
| KHV | Kinh in Ho Chi Minh City, Vietnam | EAS |
| CEU | Utah Residents (CEPH) with Northern and Western European Ancestry | EUR |
| TSI | Toscani in Italia | EUR |
| FIN | Finnish in Finland | EUR |
| GBR | British in England and Scotland | EUR |
| IBS | Iberian Population in Spain | EUR |
| YRI | Yoruba in Ibadan, Nigeria | AFR |
| LWK | Luhya in Webuye, Kenya | AFR |
| GWD | Gambian in Western Divisions in the Gambia | AFR |
| MSL | Mende in Sierra Leone | AFR |
| ESN | Esan in Nigeria | AFR |
| ASW | Americans of African Ancestry in SW USA | AFR |
| ACB | African Caribbeans in Barbados | AFR |
| MXL | Mexican Ancestry from Los Angeles USA | AMR |
| PUR | Puerto Ricans from Puerto Rico | AMR |
| CLM | Colombians from Medellin, Colombia | AMR |
| PEL | Peruvians from Lima, Peru | AMR |
| GIH | Gujarati Indian from Houston, Texas | SAS |
| PJL | Punjabi from Lahore, Pakistan | SAS |
| BEB | Bengali from Bangladesh | SAS |
| STU | Sri Lankan Tamil from the UK | SAS |
| ITU | Indian Telugu from the UK | SAS |

Source: *http://www.internationalgenome.org/category/population/*

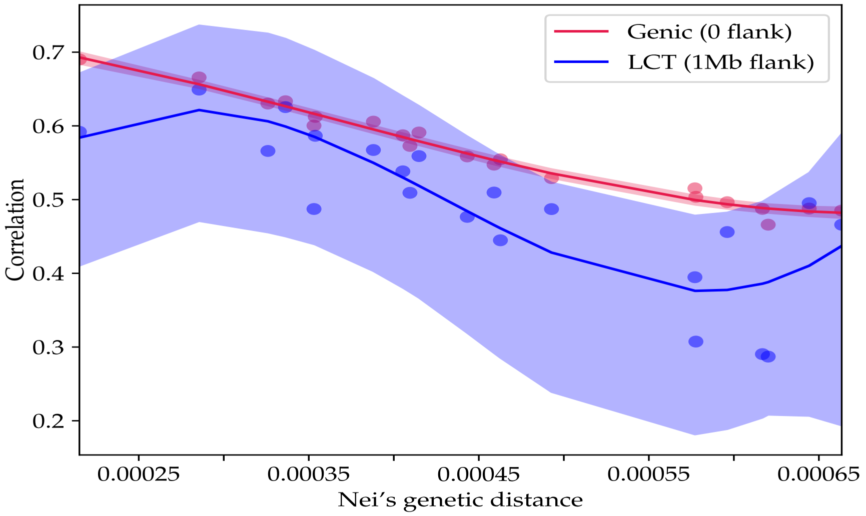


Figure 4: Correlation of |iHS| scores around LCT in African samples. Unsigned correlations either fall below genic regions taken as a whole, or they do not differ significantly from genic regions taken as a whole.

Table 2: ISI values for each population comparison in the glycophorin cluster.

| | CHB | JPT | CHS | CDX | KHV | CEU | TSI | FIN | GBR | IBS | YRI | LWK | GWD |
|---|---|---|---|---|---|---|---|---|---|---|---|---|---|
| CHB | 0.00E+00 | 9.37E-01 | 3.23E-01 | 1.05E+00 | 2.03E+00 | 1.41E+01 | 3.44E+00 | 1.04E+01 | 1.34E+01 | 1.29E+01 | 2.62E+00 | 1.67E+00 | 1.58E+00 |
| JPT | 9.37E-01 | 0.00E+00 | 7.20E-01 | 7.49E-01 | 4.62E-01 | 6.28E+00 | 3.67E+00 | 5.32E+00 | 6.40E+00 | 5.04E+00 | 1.32E+00 | 1.57E+00 | 1.88E+00 |
| CHS | 3.23E-01 | 7.20E-01 | 0.00E+00 | 4.81E-01 | 1.39E+00 | 1.99E+01 | 4.03E+00 | 1.51E+01 | 1.99E+01 | 1.82E+01 | 1.61E+00 | 1.44E+00 | 1.27E+00 |
| CDX | 1.05E+00 | 7.49E-01 | 4.81E-01 | 0.00E+00 | 1.14E+00 | 1.40E+01 | 2.85E+00 | 1.06E+01 | 1.38E+01 | 1.28E+01 | 3.08E+00 | 2.46E+00 | 2.23E+00 |
| KHV | 2.03E+00 | 4.62E-01 | 1.39E+00 | 1.14E+00 | 0.00E+00 | 2.03E+01 | 4.35E+00 | 1.53E+01 | 2.01E+01 | 1.89E+01 | 2.93E+00 | 3.14E+00 | 3.66E+00 |
| CEU | 1.41E+01 | 6.28E+00 | 1.99E+01 | 1.40E+01 | 2.03E+01 | 0.00E+00 | 2.28E+00 | 1.53E-01 | 9.13E-02 | 3.73E-01 | 1.43E+00 | 2.77E+00 | 4.90E+00 |
| TSI | 3.44E+00 | 3.67E+00 | 4.03E+00 | 2.85E+00 | 4.35E+00 | 2.28E+00 | 0.00E+00 | 1.71E+00 | 2.09E+00 | 2.75E+00 | 2.23E+00 | 2.72E+00 | 4.76E+00 |
| FIN | 1.04E+01 | 5.32E+00 | 1.51E+01 | 1.06E+01 | 1.53E+01 | 1.53E-01 | 1.71E+00 | 0.00E+00 | 6.86E-02 | 8.49E-01 | 1.46E+00 | 2.56E+00 | 4.96E+00 |
| GBR | 1.34E+01 | 6.40E+00 | 1.99E+01 | 1.38E+01 | 2.01E+01 | 9.13E-02 | 2.09E+00 | 6.86E-02 | 0.00E+00 | 6.84E-01 | 1.74E+00 | 3.34E+00 | 5.38E+00 |
| IBS | 1.29E+01 | 5.04E+00 | 1.82E+01 | 1.28E+01 | 1.89E+01 | 3.73E-01 | 2.75E+00 | 8.49E-01 | 6.84E-01 | 0.00E+00 | 1.10E+00 | 3.09E+00 | 3.66E+00 |
| YRI | 2.62E+00 | 1.32E+00 | 1.61E+00 | 3.08E+00 | 2.93E+00 | 1.43E+00 | 2.23E+00 | 1.46E+00 | 1.74E+00 | 1.10E+00 | 0.00E+00 | 2.15E+00 | 1.32E+00 |
| LWK | 1.67E+00 | 1.57E+00 | 1.44E+00 | 2.46E+00 | 3.14E+00 | 2.77E+00 | 2.72E+00 | 2.56E+00 | 3.34E+00 | 3.09E+00 | 2.15E+00 | 0.00E+00 | 5.61E+00 |
| GWD | 1.58E+00 | 1.88E+00 | 1.27E+00 | 2.23E+00 | 3.66E+00 | 4.90E+00 | 4.76E+00 | 4.96E+00 | 5.38E+00 | 3.66E+00 | 1.32E+00 | 5.61E+00 | 0.00E+00 |
| MSL | 1.51E+00 | 1.48E+00 | 1.12E+00 | 2.07E+00 | 3.00E+00 | 1.31E+00 | 9.92E-01 | 9.99E-01 | 1.20E+00 | 1.16E+00 | 1.27E+00 | 1.21E+00 | 1.26E+00 |
| ESN | 2.23E+00 | 2.46E+00 | 1.89E+00 | 3.21E+00 | 4.96E+00 | 7.23E-01 | 1.94E+00 | 1.09E+00 | 1.29E+00 | 5.11E-01 | 1.10E+00 | 2.63E+00 | 5.25E-01 |
| GIH | 2.40E+00 | 1.44E+00 | 2.44E+00 | 2.07E+00 | 2.46E+00 | 1.94E-01 | 2.26E-01 | 1.37E-01 | 1.01E-01 | 1.84E-01 | 9.36E-01 | 9.51E-01 | 1.32E+00 |
| PJL | 2.71E+00 | 3.29E+00 | 1.63E+00 | 2.10E+00 | 2.34E+00 | 4.64E+00 | 1.01E+00 | 3.63E+00 | 4.62E+00 | 5.26E+00 | 2.21E+00 | 2.11E+00 | 4.15E+00 |
| BEB | 1.97E+01 | 8.19E+00 | 2.79E+01 | 1.97E+01 | 2.86E+01 | 6.96E-02 | 2.24E+00 | 9.70E-02 | 1.96E-01 | 3.47E-01 | 1.20E+00 | 3.66E+00 | 8.27E+00 |
| STU | 1.54E+00 | 1.08E+00 | 1.27E+00 | 1.33E+00 | 1.27E+00 | 3.62E+00 | 2.33E+00 | 3.72E+00 | 4.41E+00 | 2.18E+00 | 1.45E+00 | 2.08E+00 | 8.16E-01 |
| ITU | 7.87E+00 | 3.60E+00 | 1.06E+01 | 7.54E+00 | 1.09E+01 | 1.51E-01 | 8.96E-01 | 1.37E-01 | 8.82E-02 | 2.30E-01 | 1.43E+00 | 2.74E+00 | 3.30E+00 |

Table 2 continued.

| | MSL | ESN | GIH | PJL | BEB | STU | ITU |
|---|---|---|---|---|---|---|---|
| CHB | 1.51E+00 | 2.23E+00 | 2.40E+00 | 2.71E+00 | 1.97E+01 | 1.54E+00 | 7.87E+00 |
| JPT | 1.48E+00 | 2.46E+00 | 1.44E+00 | 3.29E+00 | 8.19E+00 | 1.08E+00 | 3.60E+00 |
| CHS | 1.12E+00 | 1.89E+00 | 2.44E+00 | 1.63E+00 | 2.79E+01 | 1.27E+00 | 1.06E+01 |
| CDX | 1.05E+00 | 7.49E-01 | 4.81E-01 | 0.00E+00 | 1.14E+00 | 1.40E+01 | 2.85E+00 |
| KHV | 2.03E+00 | 4.62E-01 | 1.39E+00 | 1.14E+00 | 0.00E+00 | 2.03E+01 | 4.35E+00 |
| CEU | 1.41E+01 | 6.28E+00 | 1.99E+01 | 1.40E+01 | 2.03E+01 | 0.00E+00 | 2.28E+00 |
| TSI | 3.44E+00 | 3.67E+00 | 4.03E+00 | 2.85E+00 | 4.35E+00 | 2.28E+00 | 0.00E+00 |
| FIN | 1.04E+01 | 5.32E+00 | 1.51E+01 | 1.06E+01 | 1.53E+01 | 1.53E-01 | 1.71E+00 |
| GBR | 1.34E+01 | 6.40E+00 | 1.99E+01 | 1.38E+01 | 2.01E+01 | 9.13E-02 | 2.09E+00 |
| IBS | 1.29E+01 | 5.04E+00 | 1.82E+01 | 1.28E+01 | 1.89E+01 | 3.73E-01 | 2.75E+00 |
| YRI | 2.62E+00 | 1.32E+00 | 1.61E+00 | 3.08E+00 | 2.93E+00 | 1.43E+00 | 2.23E+00 |
| LWK | 1.67E+00 | 1.57E+00 | 1.44E+00 | 2.46E+00 | 3.14E+00 | 2.77E+00 | 2.72E+00 |
| GWD | 1.58E+00 | 1.88E+00 | 1.27E+00 | 2.23E+00 | 3.66E+00 | 4.90E+00 | 4.76E+00 |
| MSL | 1.51E+00 | 1.48E+00 | 1.12E+00 | 2.07E+00 | 3.00E+00 | 1.31E+00 | 9.92E-01 |
| ESN | 2.23E+00 | 2.46E+00 | 1.89E+00 | 3.21E+00 | 4.96E+00 | 7.23E-01 | 1.94E+00 |
| GIH | 2.40E+00 | 1.44E+00 | 2.44E+00 | 2.07E+00 | 2.46E+00 | 1.94E-01 | 2.26E-01 |
| PJL | 2.71E+00 | 3.29E+00 | 1.63E+00 | 2.10E+00 | 2.34E+00 | 4.64E+00 | 1.01E+00 |
| BEB | 1.97E+01 | 8.19E+00 | 2.79E+01 | 1.97E+01 | 2.86E+01 | 6.96E-02 | 2.24E+00 |
| STU | 1.54E+00 | 1.08E+00 | 1.27E+00 | 1.33E+00 | 1.27E+00 | 3.62E+00 | 2.33E+00 |
| ITU | 7.87E+00 | 3.60E+00 | 1.06E+01 | 7.54E+00 | 1.09E+01 | 1.51E-01 | 8.96E-01 |
